# Direct and Indirect Entomological Efficacy of Targeted Indoor Residual Spraying against *Aedes aegypti* in Iquitos, Peru

**DOI:** 10.64898/2026.05.18.725931

**Authors:** Helvio Astete, Gissella M. Vasquez, Victor Lopez, Beyquer Zambrano, Becker Reyna, Ryan C. Moore, Amy C. Morrison, Gonzalo Vazquez-Prokopec, Ryan T. Larson

**Author notes:** These authors contributed equally to this work (Amy C. Morrison, Gonzalo Vazquez-Prokopec, Ryan T. Larson).

## Abstract

**Background:** Control of *Aedes aegypti*, the primary vector of dengue and other *Aedes*-borne viruses, is challenged by insecticide resistance, limited efficacy of existing tools and the large and widespread epidemics. Targeted Indoor Residual Spraying (TIRS), a modification of traditional indoor residual spraying focused on *Ae. aegypti* resting sites, has demonstrated promising results, yet its indirect community-wide effects remain underexplored.

**Methodology/Principal Findings:** We conducted an entomological cluster-randomized controlled trial in Iquitos, Peru, to evaluate the direct and indirect entomological impacts of TIRS using pirimiphos-methyl. Thirty clusters were randomized to receive either TIRS (15 clusters, 898 structures) or standard Ministry of Health vector control activities (15 clusters, 1,018 structures). *Aedes aegypti* indoor densities were assessed in the 45 days pre-intervention and at four time points up to 255 days post-intervention using Prokopack aspiration. Generalized linear mixed models with a negative binomial link were used to estimate incidence rate ratios (IRRs) and calculate efficacy (1–IRR) for houses that received TIRS (direct effect) and untreated houses in TIRS clusters (indirect effect). Direct efficacy reached 96% at 15 days post-spraying and remained significant (40%) at 255 days post-spraying. Indirect efficacy reached 69% at 15 days and declined to 7% by 255 days post-spraying. Despite only 57% household-level TIRS coverage, both direct and indirect impacts on *Ae. aegypti* were significant during early post-intervention surveys, and after 8 months in TIRS clusters.

**Conclusions/Significance:** TIRS provided substantial and sustained reductions in indoor *Ae. aegypti* density, including measurable indirect effects in untreated homes within intervention clusters. These findings demonstrate the entomological value of TIRS even at moderate coverage levels and highlight its potential for both preventive and reactive vector control programs and should be considered for implementation by Ministries of Health in dengue-endemic urban settings as well as by the U.S. military when deployed to tropical or subtropical locations.

## Introduction

The threat of *Aedes*-borne viruses (ABV), that cause dengue, Chikungunya and Zika, is on the rise. Transmitted primarily by *Aedes aegypti*, a mosquito that thrives in homes in tropical and subtropical regions, ABVs have increased both in burden and geographic range over the past decade [1,2]. While Zika (ZIKV) and Chikungunya (CHIKV) viruses have narrowed their distribution and force of infection in recent years after their initial global expansion, dengue virus (DENV) epidemics are becoming more common and intense [2]. CHIKV vaccines are available for travelers [3] and recently a safe licensed vaccine is being deployed for DENV in high transmission settings in which people have been exposed to at least one serotype [4]. Additionally, ABV are especially problematic for U.S. Military members whose frequent deployments may put them at higher risk of severe dengue due to increased risk of multiple DENV serotypes [5]. Service members often experience more frequent exposure to disease vectors due to time spent in austere environments participating in activities such as field exercises, or HADR/ Humanitarian assistance operations. Furthermore, future military operations in distributed maritime environments will likely limit the practicality of typically heavy and cumbersome aerial application equipment. Additionally, the U.S. Military maintains a significant presence in expeditionary bases in dengue endemic areas globally. While IRS is recommended for use in installation Emergency Vector Management Plans, IRS strategies are seldom utilized in Department of War (DoW) integrated vector management strategies for *Aedes* spp [6]. Effective and locally adapted vector control options to prevent ABVs are desperately needed for both Ministries of Health and U.S. Military preventive medicine. Fortunately, the arsenal of new tools with high potential to control *Ae. aegypti* and prevent ABVs is increasing [7,8].

One such tool is the use of residual insecticides specifically targeted to locations indoors where *Ae. aegypti* commonly rests. Targeted indoor residual spraying (TIRS) emerged as an adaptation of classic indoor residual spraying (IRS) in Australia [8] by applying insecticides to walls at heights below 1.5m, under and around furniture in rooms excluding kitchens and bathrooms [9]. When implemented in Cairns, Australia, TIRS prevented up to 86% of DENV cases during an outbreak [10]. Since that early study, the entomological impact of TIRS on *Ae. aegypti* has been researched more extensively. In Mexico, the implementation of TIRS as a preventive measure (applied prior to the beginning of the ABV transmission season) led to an overall reduction of indoor adult *Ae. aegypti* of 70% for up to 7 months [11]. Such entomological impact was also associated with a drastic shift in the mosquito population age structure towards younger nulliparous females [12]. The strong preference of *Ae. aegypti* to rest at low heights, described in multiple studies across its distribution [13–15] may help explain why TIRS has such an important and lasting direct entomological impact on *Ae. aegypti*.

The implementation of vector control interventions in urban settings face many challenges (e.g., residents unwilling to accept the intervention, unwelcoming dogs which make house entry unsafe, absence of residents at the time of visits, or visit times that are inconvenient for residents) that lead to reduced coverage of interventions and surveillance activities. In Cairns, Australia, the epidemiological impact of TIRS was only statistically significant (i.e., treated houses had less DENV cases than untreated premises) when 60% of premises around a case were treated [16]. Modeling studies also predict a reduction in intervention efficacy as insecticide coverage is reduced [17,18], a measure also empirically measured in entomological field trials [19,20]. What remains harder to estimate for most vector control approaches is whether a given intervention carries an indirect entomological impact; that killing mosquitoes in a group of houses has a population-level effect that leads to a reduction in mosquitoes in nearby untreated premises. In the context of cluster-randomized trials, such effect can be quantified by splitting the overall efficacy into direct efficacy (estimate of impact on those in an intervention cluster receiving the intervention compared to those in a control cluster) an indirect efficacy (estimate of impact on those within a treatment cluster but not receiving the intervention compared to those in a control cluster) [21] (Fig 1). Indirect effects were hypothesized to be important and potentially negative for spatial repellents, given that repelling mosquitoes from one treated house may lead them to fly into nearby houses not receiving the intervention [22,23]. Recent evidence from a trial in Kenya disproved this hypothesis and demonstrated beneficial indirect effects [24]. Direct and indirect efficacy has also been estimated for bed net users within the context of malaria transmission in sub-Saharan Africa [25] and for the focalization of IRS within high risk malaria households within villages [26]. To date no study has quantified the direct and indirect effect of TIRS, representing a critical knowledge gap that needs to be addressed to assess the operational impact of not achieving full intervention coverage.

**Figure 1.**
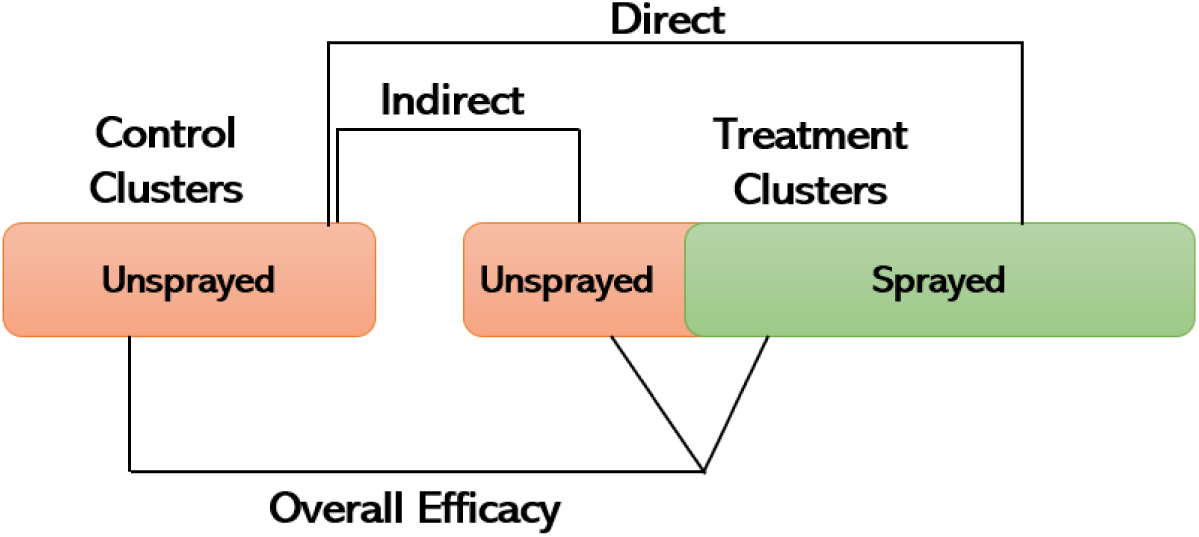
Different measures of intervention efficacy, adapted to the context of vector control interventions implemented in cluster randomized trials. The ratio between unsprayed and sprayed in the treatment clusters defines the intervention coverage. Adapted from Halloran et al. [27]

In this study, we report results from a cluster-randomized controlled trial conducted in the city of Iquitos, Peru, that evaluates the entomological impact of TIRS on *Ae. aegypti* density indoors. Our study was designed to specifically address the difference between indirect and direct entomological impact of TIRS on *Ae. aegypti*.

## Methods

### Ethical Approval

The study protocol was determined not to include human participants by the U.S. Naval Medical Research Unit SOUTH (Protocol #NAMRU6.P0007) Institutional Review Board in compliance with all U.S. Federal and Peruvian regulations governing the protection of human subjects. The protocol was reviewed and approved by the Loreto Regional Health Department, which oversees health research in Iquitos. Permission to enter households for entomological surveys and residual spray applications was obtained for every visit and after a detailed explanation on the first visit. Written information sheets describing all project activities were provided to household heads.

### Study Area

Our trial was conducted in the northern part of the Amazonian city of Iquitos, Peru, in five of 35 operational sectors designated by local vector control authorities; sectors 3-6 and sector 7 belong to the districts of Punchana and Iquitos, respectively (Fig 2). Detailed descriptions of Iquitos city have been previously published [28–30]. Housing in the study area was predominantly concrete block or brick structures with cement floors, with a smaller percentage of wood structures with dirt floors, or houses with more finished ceramic floors. Nearly all roofs in the trial were made of corrugated metal. Homes tend to be long and narrow, commonly attached to adjacent houses, without enclosed ceilings (open eaves between the top of walls), but some heterogeneity in average house size (2-500m^2^ observed across study clusters. The city has experienced both dengue and Zika virus transmission over the last 30 years [31]. Routine *Ae. aegypti* control in Iquitos consists of larviciding at ∼ 3-month intervals and health education activities utilizing billboards, radio, and TV messages focusing on dengue and its prevention. During the trial period, in response to nation-wide DENV transmission, indoor applications of Malathion using thermal foggers were conducted in all the study clusters between February 27 and March 14, 2023.

**Figure 2.**
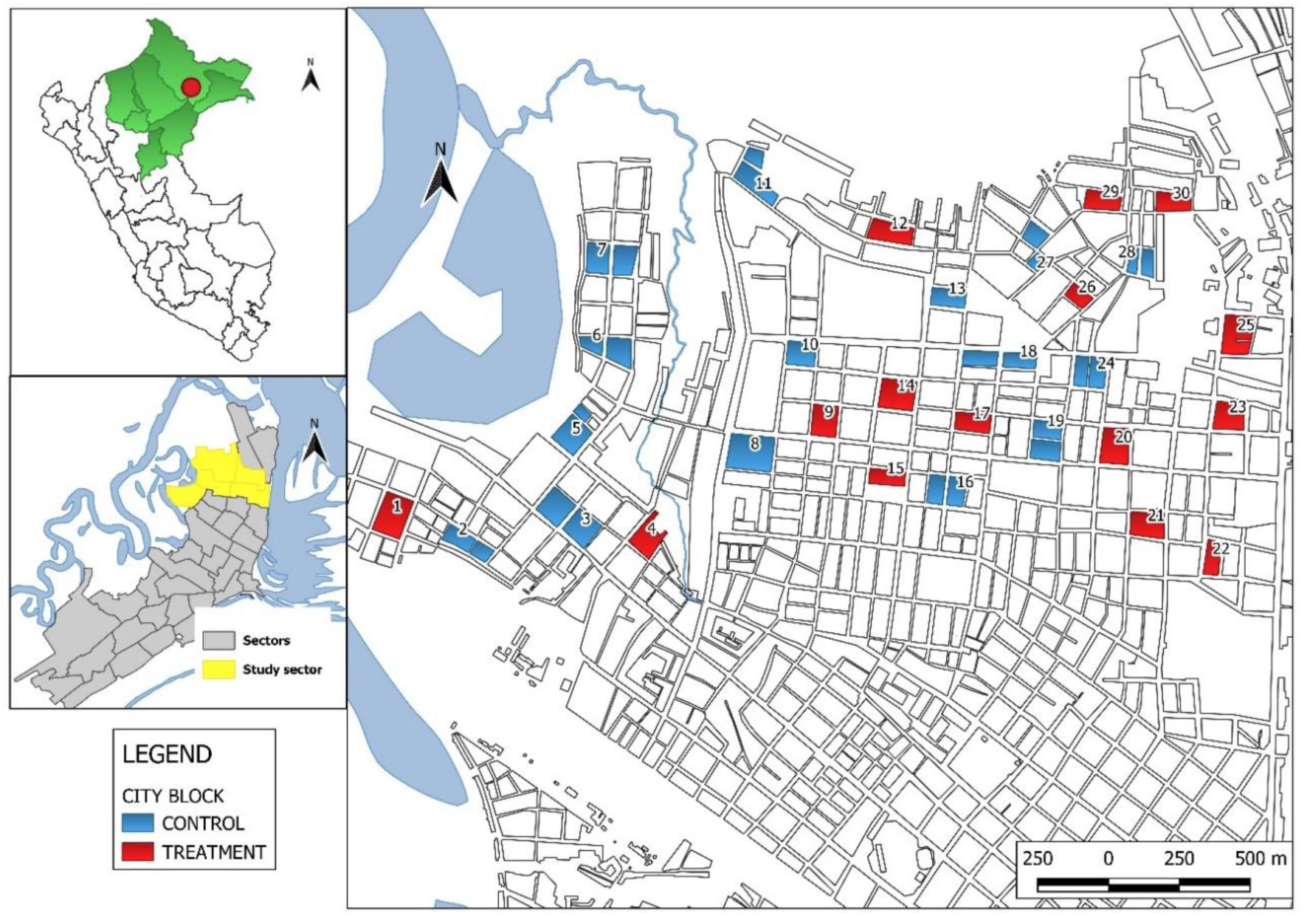
Location of study clusters (number on colored blocks) for a Targeted Indoor Residual Spray (TIRS) intervention using pirimiphos-methyl. The map was created in QGIS® v.3.34.

### Study Design

Our study was a parallel, unblinded, entomological cluster-randomized controlled vector control trial consisting of 30 clusters with 42-94 structures each. Some clusters included schools, small hotels, wearhouses, and workshops. Clusters were separated by a minimum distance of 250m, with half receiving TIRS and the other half serving as controls which received standard MINSA control measures (Fig 2). We used QGIS randomization tools to select the 15 clusters receiving TIRS. Our trial was conducted from January through November, 2023. Baseline adult mosquito surveys using Prokopack aspirators [32] (pre-TIRS) were carried out in January 2023, TIRS-applications were carried out during the second half of February 2023 with follow up Prokopack aspirations carried out in both treated and control blocks in Februray (+1-14d post application) prior to Malathion interventions by the Ministry of Health (MOH) as coordinated with our team, March (30-45d post application), July (180-195d post application), and October-November (240-255 d post application) 2023. Prokopack aspirations were carried out in all houses permitting access within a cluster during both baseline and follow up surveys, allowing us to estimate the impact of TIRS on untreated houses within the same cluster.

#### Entomological Survey Methodology

During each Prokopack aspiration survey, eight experienced collectors sampled each individual cluster simultaneously, with each collector sampling different households until all accessible houses were sampled. Each collector would obtain permission to enter the home, then proceed through all rooms (living room, dining room, bedrooms, kitchen, bathrooms, storage rooms) passing the aspirator along walls, under furniture, closets, and items on walls. Aspiration times varied with the size of the house but generally ranged from 5-10 minutes per household. Collectors would record the house number, aspiration time (minutes), number of rooms surveyed, and wall surface type (concrete, brick, plywood, wood, noting mixtures, and if painted or not). The intention of the entomology team was to survey clusters in the same order (i.e., in chronological order) throughout the study. This approach was carried in baseline, and all follow up surveys except for the first February 2023 follow up survey that coincided with MOH Malathion thermal fogging. A team member (HA) who was in constant communication with MOH personnel, switched the survey order when necessary to ensure that the first post-intervention survey was carried out prior to the MOH spray application. This ensured that the comparison between baseline and the first post-intervention survey represented an evaluation of the TIRS intervention alone but also allowed a before and after comparison for the Malathion application (between post-intervention surveys 1 and 2) in the control blocks.

#### Mosquito Processing

Adult mosquitoes were transported to the NAMRU SOUTH Iquitos laboratory, sedated at - 20°C, identified to species and sex, counted by date and house. A sample of female *Ae. aegypti* per household per collection were examined for blood meal status and scored as unfed, blood-fed (fully engorged, half-engorged, or trace amounts), or gravid [12]. These female mosquitoes were then dissected to determine their parity status (parous, nulliparous, or gravid). The number of female *Ae. aegypti* scored and dissected per household during each survey period was not consistent and depended on workload and available personnel for dissection, thus not included in data analysis.

#### TIRS Application of Pirimiphos-methyl

We applied the organophosphate insecticide pirimiphos-methyl (Actellic 300CS®) to all participating households using either Hudson pump or IK-Vector Control Super sprayers both fitted with 8002E control flow values, in each treatment cluster following WHO/PAHO protocols [33]. Briefly, both sprayers were calibrated to provide a final dose of 1 gr/m^2^, with a recommended flow rate of 580 ml/min. During the trial we documented flow rates between 580-620 ml/min for Hudson sprayers and 580 ml/min (± 5%) for IK sprayers. The insecticide was applied on exposed surfaces at a height below 1.5 m, and below and behind furniture, door, and window frames. It was not necessary to move furniture, paintings, or appliances before spraying. Per protocol, ceramic, glass, iron, or enamel paint, covered surfaces, cloths, and animal beds and anywhere in kitchens were not sprayed.

#### Analysis Plan

The primary endpoint for the trial was the density (number of *Ae. aegypti*) per house after a ∼10 min Prokopack aspiration indoors. Generalized Linear Mixed Models (GLMM) with a negative-binomial function were applied to quantify the efficacy of TIRS compared to the control arm on each sampling date. We split the entire dataset into two: a) all control houses and only houses in the intervention arm that received the intervention (Direct Efficacy set); b) all control houses and only houses in the intervention arm that did not receive the intervention (Indirect Efficacy set). We ran an overall model, including all post-intervention surveys, which included a random intercept for the survey month and another for the cluster ID. The finding of a significant overall model provided the opportunity to evaluate the efficacy of each intervention by survey date, in which case only cluster ID was used as a random intercept to nest house sub-samples to the cluster level. The direct and indirect intervention efficacy in reducing *Ae. aegypti* density was calculated as *E=1-IRR*, where IRR is the Incidence Risk Ratio calculated from each dataset using the negative-binomial GLMM [34].

## Results

A total of 8,602 *Ae. aegypti* were collected throughout the study (n=5 surveys) using Prokopack aspiration indoors. Baseline collections (prior to TIRS) included 3,223 *Ae. aegypti* (1,590 control arm and 1,633 in TIRS arm), of which 56.1% were female. Mean densities of *Ae. aegypti* per house were highest during baseline surveys ranging from 0.76 to 8.41 across the study clusters (Table 1). The mean baseline *Ae. aegypti* collected in the control and TIRS arms of the study were 2.3±1.1 and 2.9±1.9, respectively, with 95% CI values extending up to 4.2 *Ae. aegypti* adults per house (Table 1). Although there was significant heterogeneity across study clusters including the number of houses, housing and wall materials, and baseline *Ae. aegypti* per house, except for the number of households in each arm (control, 1,018; TIRS, 898) housing characteristics (Table 2) and baseline *Ae. aegypti* densities did not differ significantly between the two arms of the study (Table 3).

**Table 1.**
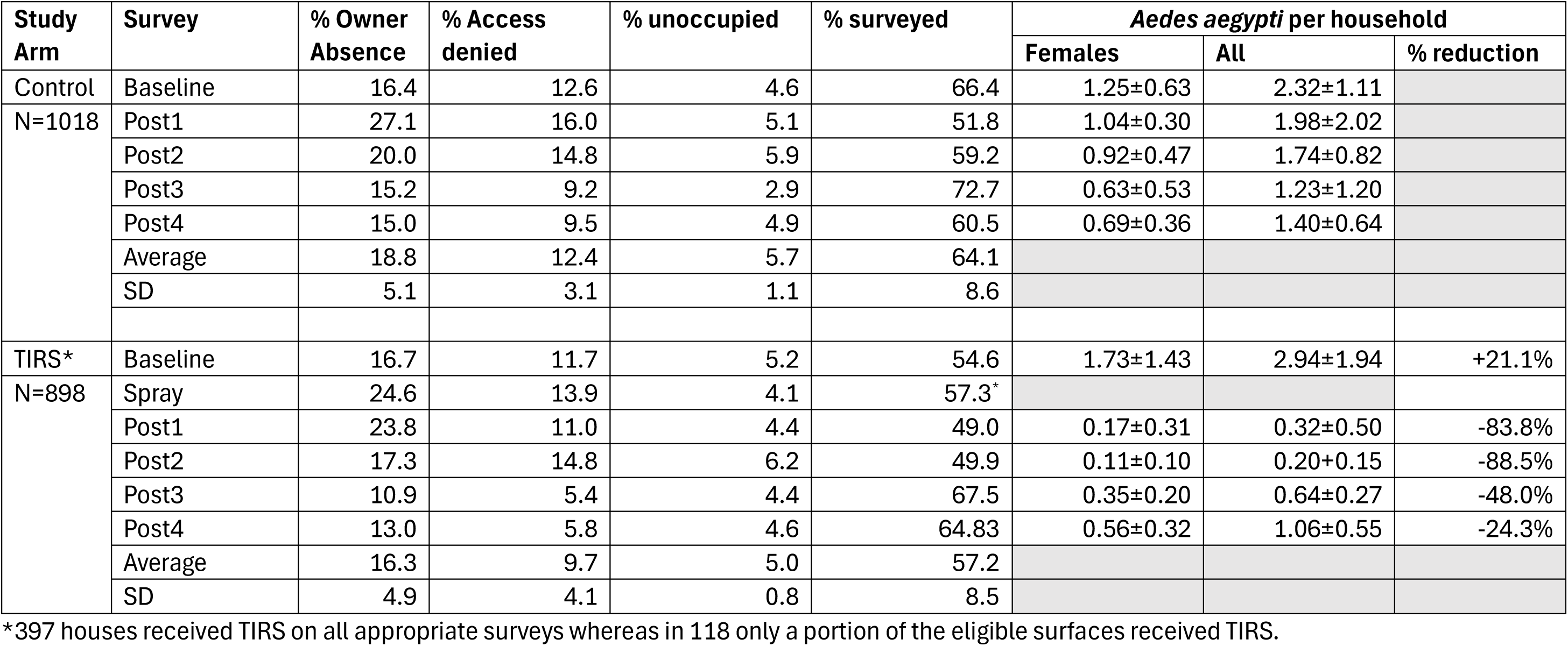
Summary of household coverage and *Aedes aegypti* densities within control and targeted indoor residual spray (TIRS) intervention areas during 2023 randomized controlled cluster trial evaluating the impact of TIRS on mosquito populations in Iquitos, Peru.

**Table 2.** Comparison of housing characteristics between control and targeted indoor residual spray (TIRS) intervention areas during 2023 randomized controlled cluster trial in Iquitos, Peru.

| Study Arm | Houses characterized | % Paint | % Concrete | % Wood | % Brick | % Plywood |
| --- | --- | --- | --- | --- | --- | --- |
| Control | 712 | 77.8±11.9 | 77.1±13.2 | 11.8±8.4 | 32.8±13.7 | 30.5±15.2 |
| TIRS | 655 | 76.2±15.6 | 78.9±13.9 | 13.1±10.7 | 29.8±11.1 | 27.3±10.9 |

**Table 3.** Results from Generalized Linear Mixed models (GLMM) testing the difference between arms (control as baseline versus TIRS) and using cluster ID as random intercept for the baseline entomological survey (prior to spraying).

| Parameter | Estimate | Std. Error | Z-value | P |
| --- | --- | --- | --- | --- |
| Intercept | 0.673 | 0.156 | 4.90 | <0.001 |
| TIRS Arm | 0.270 | 0.163 | 1.20 | 0.244 |

Out of 1,916 premises in the study area, 1,018 and 898 houses were in clusters assigned to the control arm and TIRS arm, respectively. Within the TIRS clusters, a total of 516 houses received the TIRS intervention (coverage, 57%) between 13-23 February 2023. Spraying took an average of 25.5 (SD = 14.1) minutes per house and consumed an average 0.96 (SD = 0.48) liters of insecticide per house. Not all houses in intervention areas received TIRS, with spraying coverage ranging from 43.75% to 67.8% (Table S1). Causes leading to unsprayed premises included: a) owner absence during visit (221/898 houses, 24.6%), rejection of the intervention by the household (125, 898 houses, 13.9%), uninhabited houses (37/898 houses, 4.1%). While not all premises received TIRS, 91% of houses (N= 815) received at least one entomological survey. We carried out two analyses: a) direct efficacy (including all control houses and only sprayed houses in the treatment arm) and b) indirect efficacy (all control houses and only unsprayed houses in the treatment arm).

Figure 3 shows the direct and indirect impact of TIRS on *Ae. aegypti* density indoors between one- and 255-days post-spraying, compared to the control. When looking at direct impact (Fig 3A), TIRS led to a marked reduction in *Ae. aegypti* density during the first 15 days post-intervention. When comparing control and treatment arms during the first 15 days post-TIRS, a significant reduction in adult *Ae. aegypti* density was observed (Table 4). Such significant reduction was maintained throughout all four entomological surveys, up to 255 days post-spraying (Fig 3A, Table 4). Indirect entomological impacts of TIRS were also observed (Fig 3B, Table 5), with GLMMs showing a significant reduction in untreated houses found in the treatment arm (compared to the control houses) during the first two entomological surveys, up to 45 days post-TIRS (Table 5).

**Figure 3.**
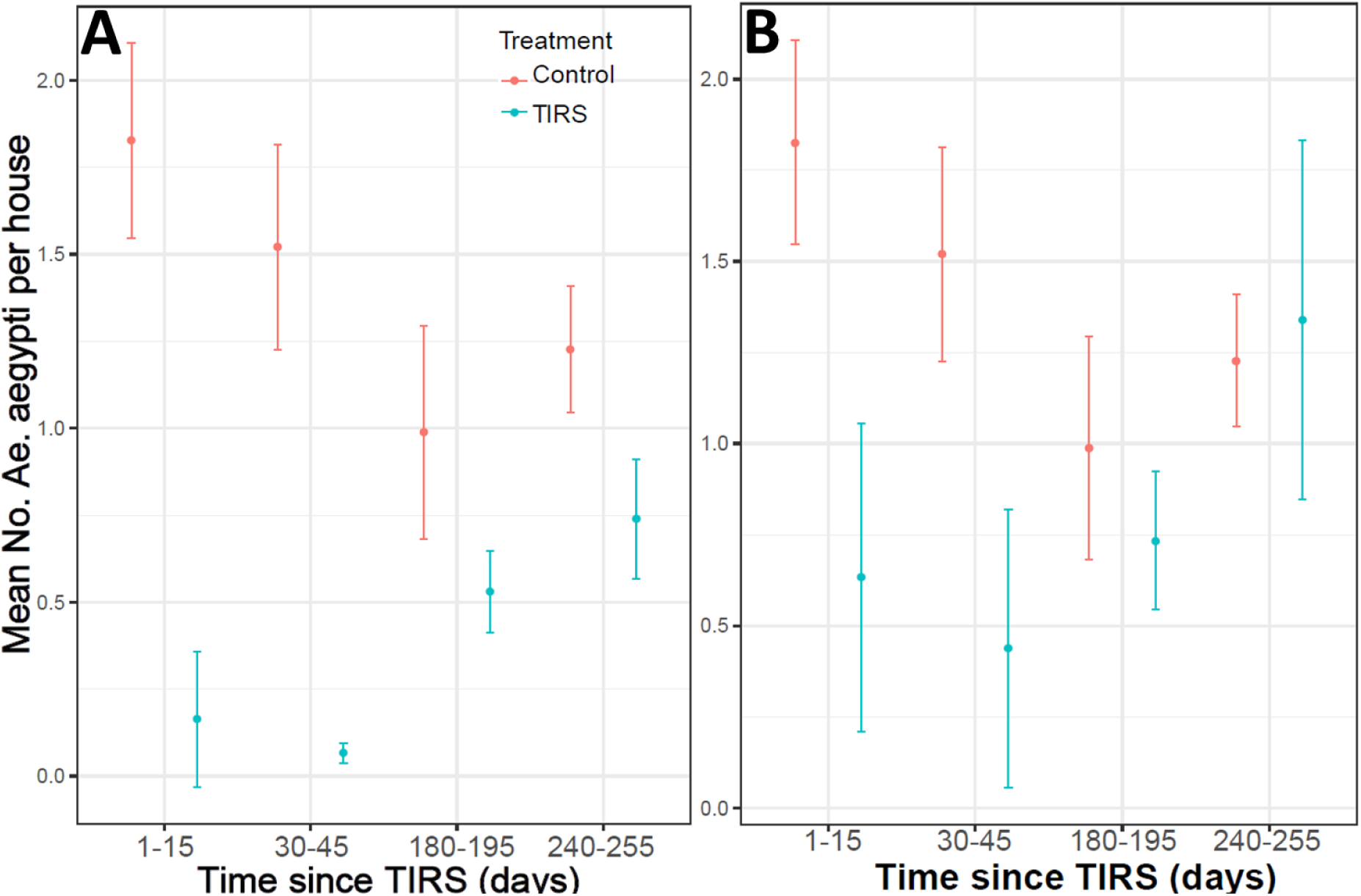
Mean density of *Ae. aegypti* adults collected with Prokopack aspirators throughout the four post-TIRS entomological surveys in houses which received the intervention (Panel A: direct efficacy dataset), intervention clusters (Panel B: indirect efficacy data set) and all control households.

**Table 4.**
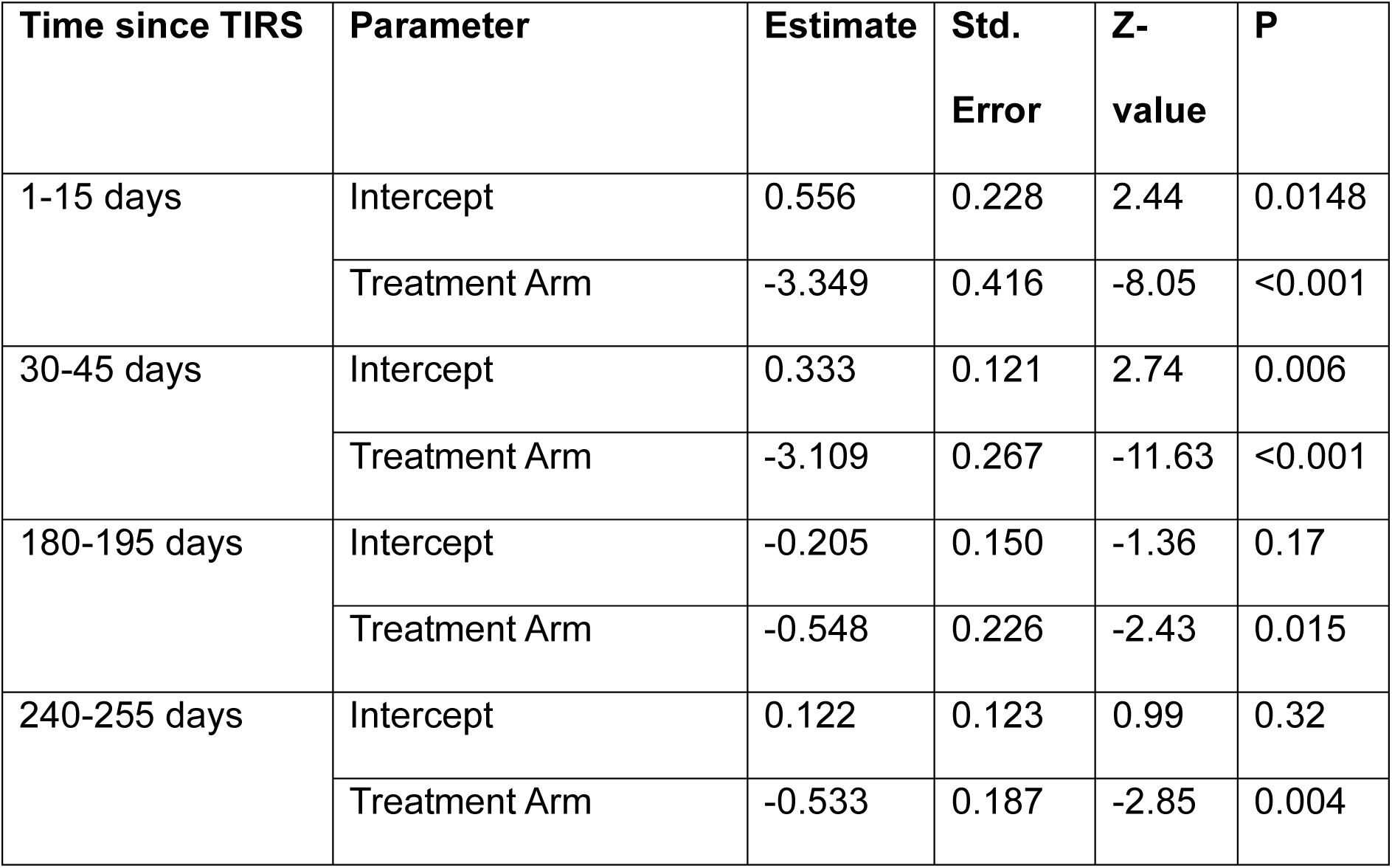
Results from Generalized Linear Mixed models (GLMM) testing the difference between arms (control as baseline versus TIRS) in the number of *Ae. aegypti* per house for the direct efficacy dataset for each of the four entomological survey conducted post-spraying. Cluster ID was used as random intercept.

| Time since TIRS | Parameter | Estimate | Std.<br>Error | Z-value | P |
| --- | --- | --- | --- | --- | --- |
| 1-15 days | Intercept | 0.556 | 0.228 | 2.44 | 0.0148 |
|  | Treatment Arm | -3.349 | 0.416 | -8.05 | <0.001 |
| 30-45 days | Intercept | 0.333 | 0.121 | 2.74 | 0.006 |
|  | Treatment Arm | -3.109 | 0.267 | -11.63 | <0.001 |
| 180-195 days | Intercept | -0.205 | 0.150 | -1.36 | 0.17 |
|  | Treatment Arm | -0.548 | 0.226 | -2.43 | 0.015 |
| 240-255 days | Intercept | 0.122 | 0.123 | 0.99 | 0.32 |
|  | Treatment Arm | -0.533 | 0.187 | -2.85 | 0.004 |

**Table 5.** Results from Generalized Linear Mixed models (GLMM) testing the difference between arms (control as baseline versus TIRS) in the number of *Ae. aegypti* per house for the indirect efficacy dataset for each of the four entomological survey conducted post-spraying. Cluster ID was used as random intercept.

| Time since TIRS | Parameter | Estimate | Std.<br>Error | Z-value | P |
| --- | --- | --- | --- | --- | --- |
| 1-15 days | Intercept | 0.570 | 0.111 | 5.11 | <0.001 |
|  | Treatment Arm | -1.179 | 0.249 | -4.73 | <0.001 |
| 30-45 days | Intercept | 1.520 | 0.050 | 30.10 | <0.001 |
|  | Treatment Arm | -1.083 | 0.078 | -14.01 | <0.001 |
| 180-195 days | Intercept | -0.207 | 0.154 | -1.34 | 0.17 |
|  | Treatment Arm | -0.234 | 0.250 | -0.94 | 0.35 |
| 240-255 days | Intercept | 0.111 | 0.147 | 0.75 | 0.45 |
|  | Treatment Arm | -0.076 | 0.241 | -0.31 | 0.75 |

Using the coefficients from the GLMM models (Table 3, Table 4) we estimated the direct and indirect efficacy of TIRS in reducing the number of *Ae. aegypti* indoors. Figure 4 shows the estimates of efficacy, using time since TIRS as a continuous variable. Direct efficacy was highest during the first 45 days post-spraying (96%) but sharply reduced as time post-TIRS increased, up to 40% at ∼200 days post-spraying (Fig 4A). Indirect efficacy followed a similar decreasing trend, but with much lower values than direct efficacy (Fig. 4B). During the first two surveys (up to 45 days), indirect efficacy ranged between 66% and 69% and sharply dropped to 21% at 180 days and to 7% at 250 days (Fig 4B).

**Figure 4.**
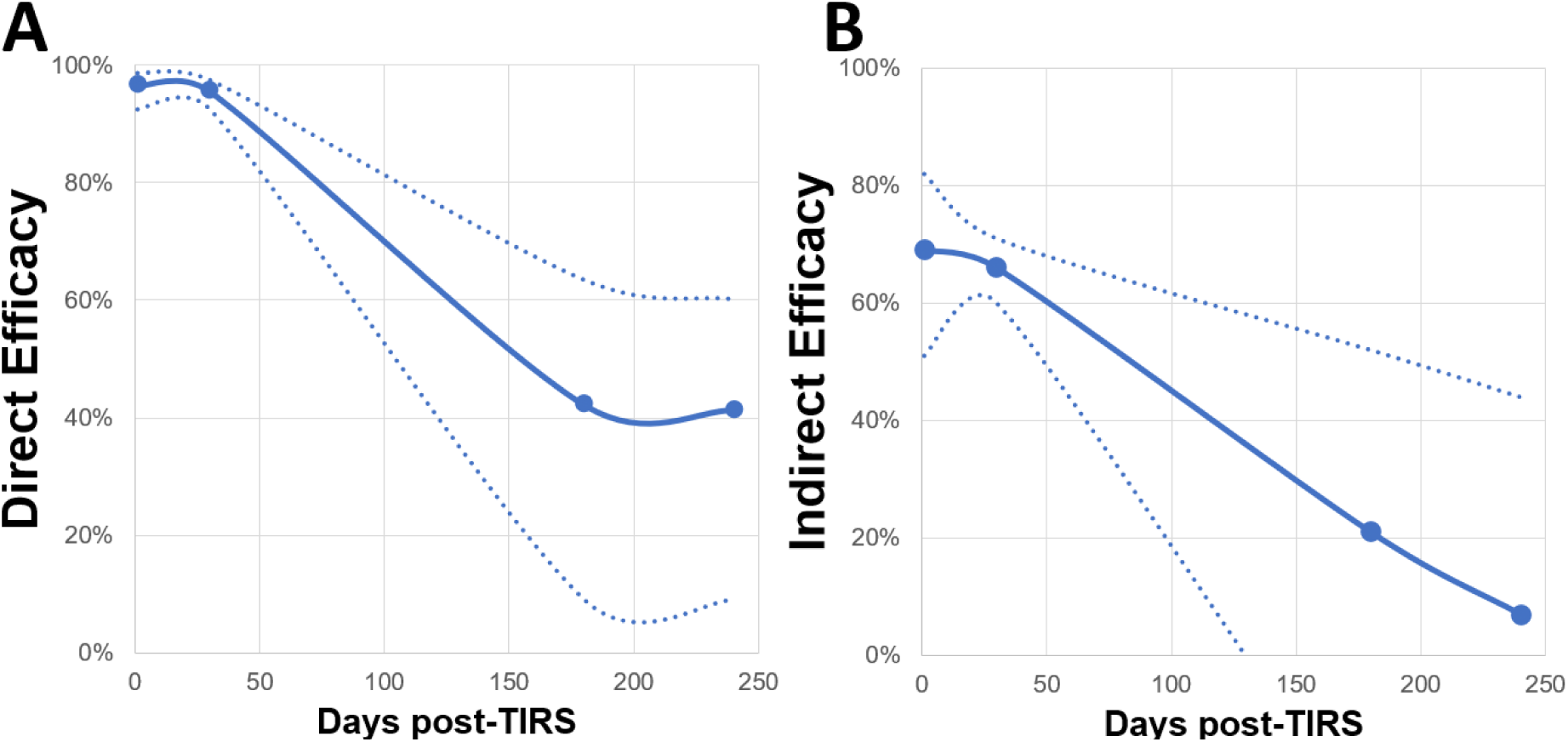
Estimates of (A) direct and (B) indirect intervention efficacy (measured as the percent reduction in number of *Ae. aegypti* indoors relative to the control) estimated from the model coefficients in Tables 3–4. Dotted lines represent the estimated binomial 95% confidence intervals.

## Discussion

The strong preference of *Ae. aegypti* for resting at low heights has been documented across different settings and housing contexts [13–15,35,36]. The modification of classic IRS to account for such *Ae. aegypti* resting behavior has led to the breakthrough development of TIRS [9], also known as selective spraying, as an innovative approach for deployment of residual insecticides within cities [33]. Prior to the conceptualization of TIRS, the use of residual insecticides for *Ae. aegypti* control was considered too costly and labor intensive for programmatic implementation. As such, most endemic countries relied heavily on vehicle-mounted space spraying or thermal fogging as the main *Ae. aegypti* control approaches, particularly during outbreaks. Evidence from multiple studies have shown that such approaches are limited in their efficacy to prevent dengue [37]. Approaches focused on the indoor environment, such as indoor space spraying using portable equipment, have been implemented by Peru and other countries as an alternative approach, with important impact on dengue prevention [38,39]. An issue with this approach is that insecticide applications have no residual effect and only lead to measurable reduction in dengue when timed right and applied on households with confirmed cases [18]. In Iquitos, applying TIRS using the organophosphate insecticide pirimiphos methyl took an average of 25 minutes (range: 5-80 min) per house (from entering house until leaving) and reduced overall *Ae. aegypti* density in treated houses between 95% at one-month post-spraying and up to 40% at ∼8 months post spraying. Such high initial effect and important residual impact open the possibility for the use of TIRS within the complex housing and epidemiological context of Iquitos.

As additional TIRS trials are completed, the body of evidence demonstrating the entomological impact of residual applications against *Ae. aegypti* continues to grow. In Mexico, using pirimiphos-methyl led to reductions in indoor adult *Ae. aegypti* densities ranging from 60% at one-month post-TIRS to 55% at 6 months [12]. Another study using two insecticide formulations found one-month efficacies of 70% (pirimiphos-methyl) and 40% (clothianidin), and 6-month efficacies of 25% (pirimiphos-methyl) and 19% (clothianidin) [11]. Such values are comparable to the direct efficacy observed in Iquitos and showcase a generalizable trend in the loss of residuality of TIRS across settings and insecticide formulations.

The very high entomological impact observed within the first 3 months post-TIRS, and extended duration of insecticidal activity for up to 6 months provide opportunity for implementation of TIRS as both preventive (applied prior to the regular arbovirus transmission season) or reactive to the occurrence of increased transmission or an outbreak context. The preventive application of TIRS in Merida, Mexico, led to the persistent reduction in indoor *Ae. aegypti* densities, which remained comparable to the densities during the low transmission season, for over a 6-month period [11]. A clinical trial [40] evaluating the epidemiological impact of the preventive application of TIRS in Mexico showed that the indoor density of *Ae. aegypti* was 59% lower in the intervention compared to control area but no significant reduction in dengue transmission in intention-to-treat analysis, although the estimated community effect of the intervention was 24% [41]. In Cairns, Australia, deploying TIRS in premises identified by contact tracing during an outbreak led to a reduction in dengue transmission by 86% [10]. In the context of Iquitos, deploying TIRS as an outbreak response method instead of indoor malathion space sprays merits evaluation. Previous studies in Iquitos have demonstrated that the short-term (1 month) dramatic reductions in *Ae. aegypti* population densities could mitigate transmission [42–44]. Area-wide space spray campaigns require high coverage rates, and three spray cycles conducted one week apart. Although spray times per house are considerably faster (∼ 2 minutes per house) than for TIRS a single 10-minute spray cycle would require similar labor requirements, and potentially less transportation costs. Added value associated with TIRS, in providing a clear and dramatic residual effect for 30 days, sufficient time to interrupt transmission cycles, as well as the clear indirect benefits to unsprayed houses which could reduce the impact of lower coverage rates suggests that TIRS may be both operationally feasible and cost-effective in comparison to street-level ULV or indoor space spraying. Future research will focus on implementation opportunities of preventive versus reactive TIRS deployment in Iquitos.

Insecticide-based interventions are dependent on the level of coverage, the residual power of insecticide formulations, and the susceptibility of vectors to the active ingredient used [19]. In Cairns, Australia, when TIRS coverage of sprayed premises around a house with a confirmed dengue case was below 60%, there was a significant increase in the number of future dengue cases [16]. Mathematical models confirmed this relationship between TIRS coverage and dengue cases [17,18,45]. One aspect rarely considered when quantifying intervention coverage is the indirect effect of an intervention on untreated premises. While lower intervention coverage may lead to treatment failure, increased coverage may protect untreated houses by killing mosquitoes that would otherwise enter in them; an analogue of ‘herd immunity’ for vaccine interventions. Applying TIRS in Iquitos led to an indirect efficacy estimate of 69% on the first week and 66% at 1-month post-spraying. Such impressive entomological reduction in untreated households provides important insight to guide intervention roll-out. In Iquitos, most unsprayed houses were closed (due to residents conducting daily activities outside their homes) rather than rejections due to safety or other concerns. Attaining very high (∼80%) coverage per block means spraying teams had to return to each block to ‘recover’ those untreated households. Pursuing a minimum coverage of 60% and advancing to cover more blocks (rather than returning to blocks to recover few households) may be more a more efficient approach for TIRS roll-out, particularly during an outbreak. The benchmark threshold of 60% TIRS coverage [16] may be useful to define minimum coverage targets.

In addition to increased adult mosquito mortality, TIRS is also known to impact *Ae. aegypti* population parameters, shifting age structure towards younger stages and leading to a non-linear reduction in survivorship compared to the control arm [12]. The combined result of such double entomological impact is a predicted reduced number of potentially infectious female *Ae. aegypti* [12]. It remains to be studied how indirect entomological impacts translate into indirect epidemiological efficacy. The link between entomological indices and dengue transmission risk is difficult and challenged by the occurrence of inapparent infections, the circulation of multiple serotypes and human mobility [46–48]. This paucity becomes more challenging when assessing the impact of vector control, given there is no identified entomological threshold that can be used to identify whether an intervention has succeeded or not [48]. What is clear is that hypothesized thresholds are very low and difficult to measure operationally, but when adult vector densities can be reduced to levels close to zero for a period extending over an extrinsic incubation period outbreaks can be mitigated significantly [44]. Mathematical models have been used to bridge this gap. Specifically for TIRS, there is ample evidence from two distinct models showing that if coverage and application of TIRS occurs in a large enough area, large reductions in dengue are expected [17,18,45]. As mentioned above, the operational cost and feasibility of TIRS for emergency control warrant investigation.

One of our studies limitations was that our study did not conduct post-spray surveys during the period 45-180 days after spray where a significant drop in efficacy was observed. A precise estimate of how many months adult densities were >80% less in treated clusters would better define the value added of using TIRS over indoor space sprays. For example, if reductions were maintained for at least three months with a single application the expected impact on virus transmission would potentially ensure that no additional interventions would be needed during a single transmission season. Understanding the duration of the indirect spray benefits is also critical to evaluate TIRS as a vector control strategy. We also recognize that the most relevant evaluation of TIRS would include evaluation of public health impact on either new dengue cases or transmission measured through seroconversion, but this was beyond the scope of our trial. It is important to note that our trial was conducted with the collaboration of the Loreto Regional Health Department who provided the pirimiphos-methyl that was purchased for malaria control and was reaching its expiration date. In many ways this was an opportunistic experiment that demonstrated the potential of the TIRS strategy. Also absent was a systematic evaluation of community acceptance and potential development of insecticide resistance within the *Ae. aegypti* population. The latter is an important consideration before proposing large-scale deployment of any kind of IRS in urban communities. We can, however, share that informal community comments were very positive towards TIRS, in large part because sprayed houses could see direct results (dead cockroaches) and reported less mosquito biting, something often less apparent with space sprays.

Although our results were very encouraging, deployment of TIRS are a larger scale requires appropriate planning, insurance of appropriate equipment that are properly calibrated, and evaluation of the cost-efficiency of the interventions. Strategies to procure and store appropriate active ingredients in coordination with programs of other vector control programs (malaria, chagas, leishmaniasis) is needed. Although local officials, after seeing preliminary data from this study, were tempted to deploy TIRS across Iquitos, like most chemical interventions, appropriate implementation is essential for success.

The public health value of TIRS has been recognized by the World Health Organization [49], yet this approach has not been widely implemented. PAHO has recognized this major gap and has been instrumental in the development of tools and frameworks for the adoption of TIRS in the Americas [33]. At the core of the PAHO framework is the use of historical dengue case data to identify transmission hotspots within which TIRS and other approaches such as Wolbachia could be more cost-effectively deployed. which is embedded within a regional plan for implementing interventions based on epidemiological scenarios and the identification of transmission hot-spots [50]. Such effort has led to recent major progress. In early 2025, the Ministry of Health of Brazil adopted the programmatic deployment of TIRS in areas of high potential for human-mosquito contacts such as households, premises with high numbers of people such as health posts, elderly centers, recycling centers, bus terminals, universities and schools located within hot-spot areas[51]. Similarly, on January 2025 Mexico launched its National Plan for the Control of Dengue and other arboviral diseases 2025-2030 in which one of the tools would be TIRS applied within hot-spot areas [52]. Information from studies such as ours will provide important insight to decision makers in other countries such as Peru about the value of TIRS and, ultimately, its ability to reduce the burden of dengue and other *Aedes*-borne diseases.

The use of TIRS could also be adapted for use in military operations in tropical or subtropical locations. Despite the U.S. military’s emphasis on skin repellents and permethrin treated uniforms, dengue ranks among the most commonly reported vector borne diseases detected in U.S. members [5,53]. Thus, new control methods are also needed to protect U.S. members in dengue endemic areas. The current study demonstrates that the longer lasting TIRS may be a more practical application in deployed military environments where trained pesticides applicators are often limited; this approach would require less frequent visits of the applicators to remote outposts for pesticide applications in comparison to space sprays. While several studies have demonstrated the long-lasting effectiveness of TIRS in civilian communities, future studies should evaluate the efficacy of residual insecticides on materials likely to be used in military operations for shelter such as tents. Existing IPMP’s and EVMP’s at established Expeditionary bases may also be augmented with TIRS enhancing Force Health protection in endemic areas where DoD maintains forces of readiness.

## Acknowledgments

We are grateful to the Ministerio de Agricultura y Riego de Peru, Dirección General Forestal y de Fauna Silvestre for permission to conduct these studies under the auspices of Resolución Directoral No. 132-2019-MINAGRI-SERFOR-DGGSPFFS. We thank the residents of Iquitos, Peru, for allowing us to undertake this study in and around their homes. We greatly appreciate the support of the Loreto Regional Health Department including Gloria Diaz, Raul Pinedo, and Jose Gonzales, who provided both pirimiphos-methyl (Actellic 300CS®). We are grateful to the Dirección General de Salud Ambiental (DIGESA) fumigation team for their support in the field. We thank Karin Escobedo who provided support for insecticide resistance testing, and Luz Romero and Maria Bosantes who assisted with data entry. We are grateful to Jhon Bardales Cardenas, Cesar Campos Cardenas, John Campos, Guillermo Elespuru Hidalgo, Wilmer Murayari, Rene Pinedo, Edward Sifuentes, and Eder Torres, the field team who carried out the entomological surveys. We thank Pablo Manrique-Saide and Azael Che-Mendoza who provided training in TIRS to the NAMRU SOUTH team leader.

## Funding

This study was funded by the DHP Restoral Program. ACM received salary support from the U.S. National Institute of Allergy and Infectious Diseases (NIH/NIAID) award numbers U01AI151814, R01AI176380, and R01AI182408. The funders had no role in study design, data collection and analysis, decision to publish, or preparation of the manuscript.

## Disclaimer

The views expressed in this article reflect the results of research conducted by the authors and do not necessarily reflect the official policy or position of the Department of the Navy, Department of Defense, nor the U.S. Government.

## Copyright statement

Some authors of this manuscript are military service members or employees of the U.S. Government. This work was prepared as part of their official duties. Title 17 U.S.C. §105 provides that Copyright protection under this Title is not available for any work of the United States Government. Title 17 U.S.C. §101 defines a U.S. Government work as a work prepared by a military Service member or employee of the U.S. Government as part of that person’s official duties.

## Data availability

All data used in this manuscript is available at Vazquez-Prokopec, Gonzalo (2026), “Direct and Indirect Entomological Efficacy of Targeted Indoor Residual Spraying in Iquitos, Peru”, Mendeley Data, V1, doi: 10.17632/vwfv3s28vc.1

## Notes

### Competing Interest Statement

The authors have declared no competing interest.

